# Genetic and molecular mechanism for distinct clinical phenotypes conveyed by allelic truncating mutations implicated in *FBN1*

**DOI:** 10.1101/726646

**Authors:** Mao Lin, Zhenlei Liu, Gang Liu, Sen Zhao, Chao Li, Weisheng Chen, Zeynep Coban Akdemir, Jiachen Lin, Xiaofei Song, Shengru Wang, Qiming Xu, Yanxue Zhao, Lianlei Wang, Yuanqiang Zhang, Zihui Yan, Sen Liu, Jiaqi Liu, Yixin Chen, Xu Yang, Tianshu Sun, Xin-Zhuang Yang, Yuchen Niu, Xiaoxin Li, Wesley You, Bintao Qiu, Chen Ding, Pengfei Liu, Shuyang Zhang, Claudia M. B. Carvalho, Jennifer E. Posey, Guixing Qiu, James R. Lupski, Zhihong Wu, Jianguo Zhang, Nan Wu, on behalf of the Deciphering Disorders Involving Scoliosis and COmorbidities (DISCO) study

**Author notes:** Corresponding authors: Nan Wu, MD,; Jianguo Zhang MD,; Zhihong Wu, MD,. These authors have contributed equally to this study.

## Abstract

The molecular and genetic mechanisms by which different single nucleotide variant (SNV) alleles in specific genes, or at the same genetic locus, bring about distinct disease phenotypes often remain unclear. Allelic truncating mutations of *fibrillin-1(FBN1)* cause either classical Marfan syndrome (MFS) or a more severe phenotype associated with Marfanoid-progeroid-lipodystrophy syndrome (MPLS). A total of three Marfan syndrome/Marfanoid patients (2 singletons and 1 parent-offspring trio) were recruited. Targeted next-generation sequencing was performed on all the participants. We analyzed the molecular diagnosis, patient clinical features, and the potential molecular mechanism involved in the MPLS subject in our cohort. We investigated a small cohort, consisting of two classical MFS and one MPLS patient from China, whose clinical presentation included scoliosis potentially requiring surgical intervention. We provide evidence that most nonsense and frameshift mutations lead to *FBN1* null alleles due to mutant mRNA transcript degradation. In contrast, the more severe disease phenotype, MPLS, is caused by mutant mRNAs that are predicted to escape the nonsense mediated decay (NMD) surveillance pathway, making a mutant protein that exerts a dominant negative interference effect to *FBN1* thus generating a gain-of-function (GoF) rather than a loss-of-function (LoF) allele as in MFS. Overall, we provide direct evidence that a dominant negative interaction of *FBN1* potentially explains the distinct clinical phenotype in MPLS patients through genetic and functional analysis of the first Chinese patient with MPLS. Moreover, our study expands the mutation spectrum of *FBN1* and highlights the potential molecular mechanism for MPLS patients.

---

Marfan syndrome (MFS; MIM: #154700) refers to a heritable autosomal dominant disease trait of fibrous connective tissue due to heterozygous mutations in the fibrillin-1 gene (*FBN1*; 134797) on chromosome 15q21. The cardinal phenotypic features allowing for clinical diagnosis primarily occur in the skeletal, ocular, and cardiovascular systems (DIETZ 2015). MFS is manifest by clinical features/findings involving the skeletal (tall stature, disproportionately long limbs and digits [arachnodactyly], anterior chest wall deformity, mild to moderate joint laxity and frequent spinal deformity [especially scoliosis], cardiovascular (increased risk for aortic root dilation and/or dissection), and ocular (*ectopia lentis*) systems (LOEYS *et al*. 2010). Marfanoid-progeroid-lipodystrophy syndrome (MPLS; MIM: #616914) is a more recently-clarified fibrillinopathy, and also a complex disease characterized by accelerated aging and postnatal lipodystrophy, poor postnatal weight gain and characteristic dysmorphic facial features that have very rarely been clinically recognized and reported (GOLDBLATT *et al*. 2011; PASSARGE *et al*. 2016). Recent studies have implicated a potential hormone, named asprosin, encoded by the *FBN1* locus as a mediator of the lipodystrophy phenotype (ROMERE *et al*. 2016; DUERRSCHMID *et al*. 2017). All previously reported MPLS individuals consistently harbor heterozygous truncating mutations in exon 64, which leads to premature stop codons in the C-terminal domain of *FBN1* (GOLDBLATT *et al*. 2011; SONG *et al*. 2012; GARG AND XING 2014; JACQUINET *et al*. 2014; PASSARGE *et al*. 2016; ROMERE *et al*. 2016).

In this study, we present clinical and genetic, genomic and molecular data from three unrelated subjects with Marfan/Marfanoid syndrome, including the first case of clinically recognized MPLS in the Chinese population. The potential effects of truncating mutations of *FBN1* which are implicated in MPLS subjects conveying distinct dominant negative alleles and clinically distinguishable autosomal dominant (AD) disease traits were ascertained with functional assays.

## Materials and methods

### Participant recruitment and Sample preparation

A total of three Marfan syndrome/Marfanoid patients (2 singletons and 1 parent-offspring trio) were consecutively recruited through the Deciphering Disorders Involving Scoliosis & Comorbidities (DISCO) project (www.discostudy.org) at Peking Union Medical College Hospital (PUMCH). Written informed consent was provided by patients or their parents. Peripheral blood samples were extracted from affected probands and corresponding unaffected parents if available. Genomic DNA was extracted with the QIAamp DNA Blood Mini Kit (QIAGEN, Germany) according to the manufacturer’s instructions.

### Targeted next-generation sequencing, Variant processing, filtering and annotation

To establish an NGS panel maximally covering known genes implicated in the cause of Mendelian diseases with vertebral malformation phenotypes, we performed a systematic literature and database search for congenital scoliosis (CS) related disease. After manual review, a total of 344 genes were found, corresponding to 457 CS related monogenic phenotypes. Also included was an additional set of 220 genes whose protein products are involved in the pathways related to vertebral development or in which pathogenic variant alleles have been reported in association with CS phenotypes in animal models, but not yet linked to human diseases with CS phenotypes. The target panel set comprises 564 genes, 6458 exons and flanking 30bp regions, and 2.97 Mb in total genomic size. DNA Probes for target capture were designed and purchased from NimbleGen (Roche, Switzerland) (https://design.nimblegen.com/nimbledesign/app). The final set of DNA probes was expected to cover 99.7% of the targeted regions. In addition to the previously published *TBX6* gene featuring a compound heterozygous inheritance model (WU *et al*. 2015), the *FBN1* gene was also targeted in the panel. A targeted sequence enrichment library was prepared following a previously described protocol (ASAN *et al*. 2011). For each sample, 1 μg of genomic DNA was used as starting material. Genomic DNA was fragmented to a size of ∼200-250 bp. The fragmented DNA was end-repaired, capped with A-tailing and subject to ligation of indexed adaptors. After 5 cycles of PCR amplification, each indexed product was pooled and hybridized with DNA probes for targeted capture in one solution capture reaction. The product of targeted enrichment DNA was further subject to 14 cycles of PCR amplification, followed by final library yield validation by Bioanalyzer analysis (Agilent, USA) and qPCR quantification. Sequencing was performed with the PE100 mode on an Illumina Hiseq2500 sequencer (Illumina, CA, USA).

After generating raw sequence data, a perl script was used to remove low-quality and adaptor-contaminated reads. The remaining reads were mapped onto human reference genome assembly hg19 (GRCh37, http://genome.ucsc.edu/) with the BWA mapper. Additionally, single nucleotide variants (SNVs) and small insertions and deletions (indels) were called with the GATK (v2.2-3) bioinformatics package. Variants were annotated using an in-house developed annotation pipeline. We filtered out variants with common and high frequencies (Minor allele frequency >1%) in the public databases (the 1000 Genomes Project, the Exome Sequencing Project 6500 (http://evs.gs.washington.edu/EVS/), ExAC (http://exac.broadinstitute.org) or an in-house WES control dataset) that were also not functionally relevant (deep intronic (>30bp), untranslated regions, or synonymous SNVs, or noncoding indels). We also annotated the detected variants using a customized database based on the Human Gene Mutation Database (HGMD) and Online Mendelian Inheritance in Man (OMIM) (https://omim.org/).

### Sanger sequencing

Genomic DNA from all individuals was subjected to Sanger sequencing for orthogonal confirmation of all identified variants.

### Prediction of probability of NMD events

For allelic truncating frameshift variants identified in our cohort and previously reported, *in-silico* predictions of NMD events were implemented using the online NMDEscPredictor (COBAN-AKDEMIR *et al*. 2018).

### Plasmid construction and mutagenesis

A cDNA fragment containing full-length *FBN1* (GenBank ID: NM_000138.4) cDNA derived from human muscle and having suitable restriction sites was PCR-amplified using KOD-Plus-Neo (Toyobo, Japan). The PCR amplicons were cloned into the *Nhe*I and *Sac*II sites of the pEGFP-N1 expression vector (Clontech, Takara Bio, Japan). Mutant EGFP-FBN1 plasmids were generated by site-directed mutagenesis and their construction confirmed by direct Sanger dideoxy sequencing. G2003R which was previously reported in an adolescent idiopathic scoliosis (AIS) case was used as a positive control (BUCHAN *et al*. 2014). Mutations of Tyr2596Thrfs*86, Glu2759Cysfs*9 and G2003R were introduced into a wild-type (WT) pEGFP-FBN1 QuikChange Lightning Site-directed Mutagenesis Kit (Agilent Technologies, CA, USA) according to the manufacturer’s instructions. The resulting three mutant plasmids pEGFP-FBN1-Tyr2596Thrfs*86, pEGFP-FBN1-Glu2759Cysfs*9 and pEGFP-FBN1-Gly2003Arg were used for functional studies.

### Cell culture and transfection

HEK293T cells were cultured in DMEM (Gibco, Waltham, MA, USA) + 10% FCS (Biological Industries, Cromwell, USA) + 1% penicillin/streptomycin (Gibco-Life Technologies, USA) at 37°C with 5% CO2. Cells were transfected/co-transfected with a mutant and/or a WT pEGFP-FBN1 using Lipofectamine3000 (Invitrogen, CA, USA) according to the manufacturers’ instructions. Six hours after transfection, the medium was replaced with fresh complete DMEM culture medium, and the cells were further incubated for 48 h. Total proteins were extracted and analyzed by Western blot and SDD-AGE, respectively.

### Western blot

Cells were lysed with modified RIPA (50 mM Tris-HCL, 1% NP40, 0.25% Na-deoxycholate, 150 mM NaCl, and 1 mM EDTA; Complete^TM^ Protease Inhibitor Cocktail [Roche]), and protein concentrations were determined with the BCA-Kit (Pierce). A total amount of 5 mg protein was size separated on an 8% SDS polyacrylamide gel, and proteins were electrophoresed and transferred to nitrocellulose membranes. Membranes were blocked in powdered milk for 30 min at room temperature (RT, 25-degrees Celsius), and primary antibodies (Phospho-Smad2 (Ser245/250/255) Antibody, Cell Signaling Technology; anti-GAPDH, Millipore) were incubated overnight at 4°C. After washing, the corresponding horseradish-peroxidase-coupled goat anti-rabbit secondary antibodies (KPL) were incubated for 1h at RT. Bands were visualized with the WesternBright ECL chemiluminescence system (Advansta, CA, USA). This experiment was performed three times with different cell lysates. Chemiluminescent signals were quantified using ImageJ (SCHNEIDER *et al*. 2012).

### SDD-AGE

For Semi-Denaturing Detergent-Agarose Gel Electrophoresis (SDD-AGE) (BERCHOWITZ *et al*. 2015), HEK293T cells were co-transfected with wildtype (pEGFP-FBN1) and mutants (pEGFP-FBN1-Tyr2596Thrfs*86; pEGFP-FBN1-Glu2759Cysfs*9) for 6 h, and then incubated for 48 h. After 48 h incubation at 37°C with 5% CO2 in the thermotank, cells were harvested by centrifugation at 3000 rcf for 2 min, resuspension in 200 μL water, and subsequent centrifugation. Approximately 100 μl of acid-washed cells were then added to each well followed by 120 μL lysis buffer (100 mM Tris pH 8, 1% Triton X-100, 50 mM β-mercaptoethanol, 3% HALT protease inhibitor cocktail, 30 mM N-ethylmaleimide, and 12.5 U/mL Benzonase nuclease). Blocks were then sealed with a rubber mat (Nunc 276002) and shaken at max speed two times for 3 min on a Qiagen Tissuelyzer 2. To each well was then added 35 μL 4X sample buffer (2X TAE, 20% glycerol, 8% SDS, 0.01% bromophenol blue). The blocks were then vortexed briefly and allowed to incubate at RT for three minutes, followed by centrifugation for 2 min at 3000 rcf to remove cell debris. Electrophoresis and capillary blotting to Hybond ECL nitrocellulose were performed as described. Proteins were transferred to a polyvinylidene difluoride membrane and probed with well-characterized monoclonal antibody of anti-Fibrillin 1 (Abcam ab124334, Cambridge, UK).

### Statistical analysis

SPSS software, version 17.0 (IBM Corporation; USA), was used to conduct student’s two-tailed t-test comparing values of test and control samples. *P*-values of less than 0.05 were considered statistically significant.

### Data availability statement

All reagents and plasmids are available upon request. Table S1 and S2 contain a list and descriptions of summary statistics of targeted genes for CS panel and targeted sequencing of 564 genes, respectively. Supplemental files have been submitted to figshare. All mutations identified in this study have been submitted to the Clinvar database (Accession ID: VCV000617941, VCV000617942, VCV000617943).

## Results

### Clinical characterization and novel allelic variants of individuals with MFS/Marfanoid disease

#### Subject XH253

The proband was a 10-year-old Chinese girl (**Fig.1A-D**) who presented with kyphoscoliosis confirmed by radiologic imaging. She was referred to our orthopedic spine specialist for further evaluation and management. Whole-spine X ray examination revealed scoliosis with three curves. Cobb angle of the major curvature was 43 degrees, and she displayed a flat back and thoracolumbar kyphosis. Transthoracic echocardiographic evaluation revealed moderate mitral valve insufficiency and an aortic root measurement of 3.6 cm (Z-score=4.5 when standardized to age and body surface area). Skeletal system abnormalities such as long and thin limbs, and arachnodactyly were observed in the proband. Diminution of vision (myopia) occurred when she was 5 years old. The proband fulfilled the clinical criteria for classical MFS according to Ghent classification (**Table.1**), with a total of 7 points for the score in the revised Ghent Criteria (LOEYS *et al*. 2010).A heterozygous frameshift deletion c.7785delC (p.Tyr2596Thrfs*86) was identified in *FBN1* (RefSeq transcript number: NM_000138.4) in the proband via targeted NGS, and subsequently confirmed by further validation using an orthogonal experimental approach of Sanger sequencing (**Fig.1E**).

**Figure 1.**
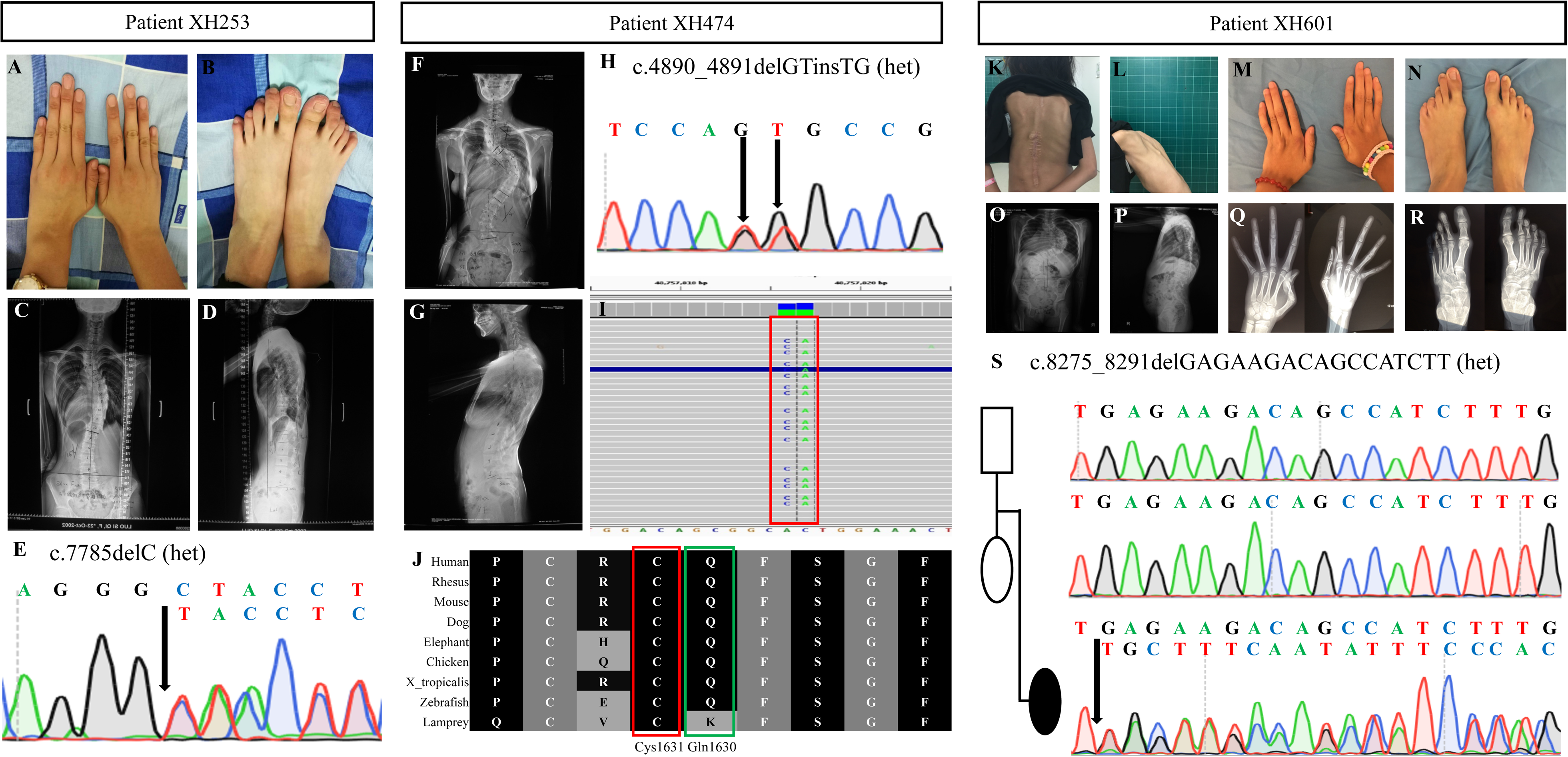
Clinical and genetic manifestations of the research subjects (A, B) Pictures of patient XH253 indicated bilateral arachnodactyly of hands and feet. (C, D) The anterior and lateral view of Patient XH253 demonstrated scoliosis on whole-spine X-ray images.(E) Sanger sequencing of XH253 verified a novel heterozygous frameshift variant of p.Tyr2596Thrfs*86 in *FBN1*. (F, G) The anterior and lateral view of subject XH474 demonstrated scoliosis on whole-spine X-ray images. (H) Sanger sequencing verified monoallelic missense mutations of p.Gln1630His and p.Cys1631Gly in *FBN1*. (I) Integrative Genomics Viewer (IGV) displays varying level of alignment reads detail depending on the zoom level and uses red box and transparency to highlight monoallelic missense variants in the exome data from XH474. (J) Conservation analysis of amino acid residues of *FBN1* among vertebrates. Lines indicate homologous amino acid sequences in selected vertebrates. Note the strong conservation of Cys1631 among vertebrates highlighted in red and Gln1630 highlighted in green, respectively. (K, L, M, N) Photographic images of subject XH601 who presented with scoliosis, short stature and subcutaneous fat reduction and arachnodactyly anomalies of hands and feet.(O, P) The anterior and lateral view demonstrated severe scoliosis on whole-spine X-ray images. (Q, R) Hand and foot radiographs showed bilateral metacarpophalangeal dislocation, interosseous atrophy, claw hands and dolichostenomelia. (S) Sanger sequencing of XH601 parent-offspring trio verified a novel de novo heterozygous frameshift mutation of p.Glu2759Cysfs*9 in *FBN1*. The RefSeq transcript number of *FBN1* is NM_000138.4.

**Table 1.**
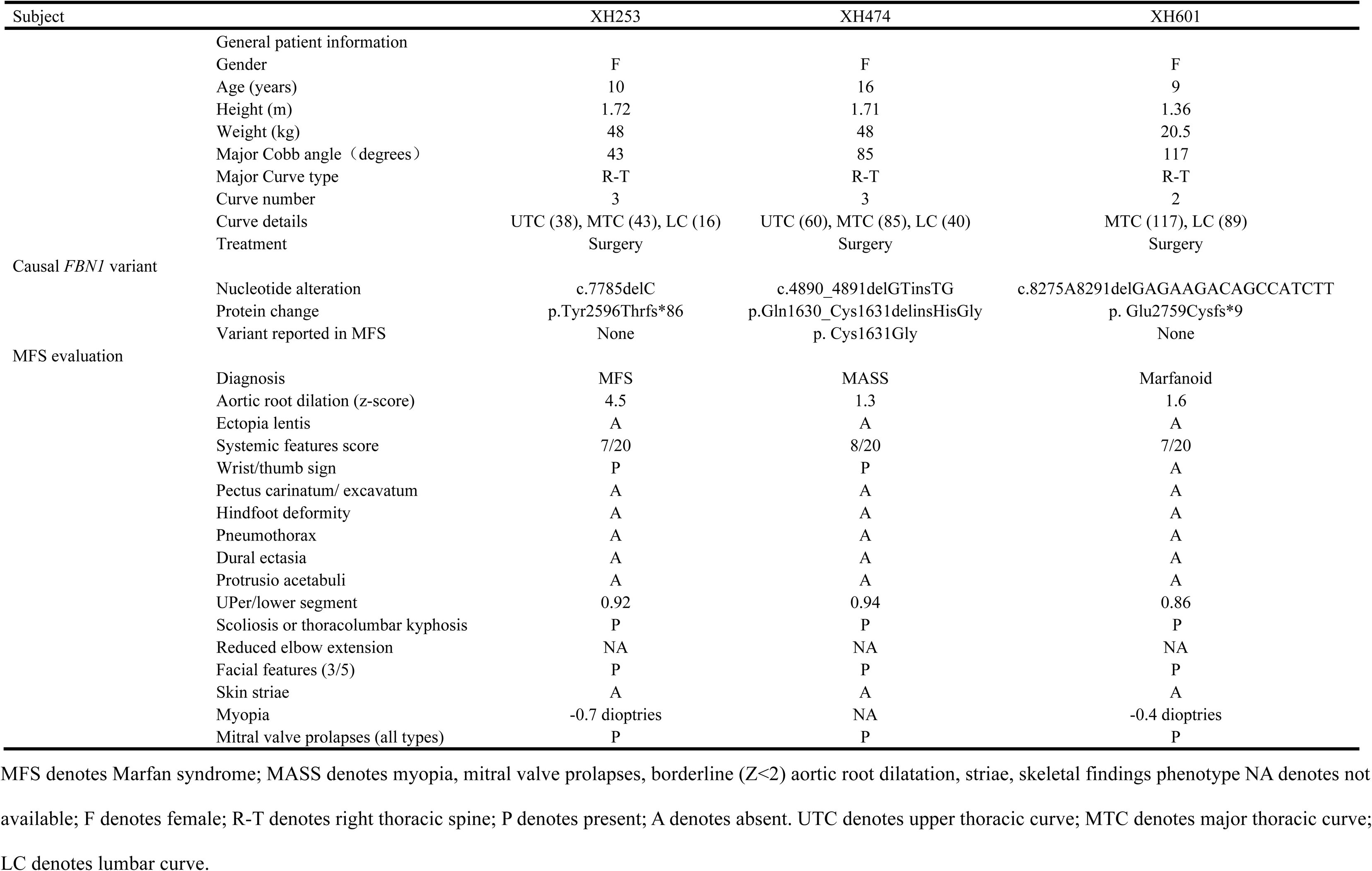
Clinical features of Marfan syndrome patients re-evaluated for according to the revised Ghent nosology

#### Subject XH474

The proband was a 16-year-old Chinese girl (**Fig.1F, G**) who presented with *pectus excavatum* and scoliosis confirmed by radiologic imaging. Transthoracic echocardiographic evaluation revealed anterior mitral leaflet prolapse and moderate mitral valve insufficiency and an aortic root diameter measurement at the level of the sinuses of Valsalva of 2.6 cm (Z-score=1.3 when standardized to age and body surface area). Skeletal system abnormalities were present, such as tall stature with a height of 171 cm (P_97_; +3SD), slender body habitus with a weight of 48 kg (P_25_, -1SD), and very thin and long upper and lower limbs. Both of her hands indicated an appearance of arachnodactyly; wrist sign and thumb sign were positive. According to the revised Ghent nosology in 2010, an *FBN1* mutation is identified in this sporadic case but aortic root measurements are still below Z score=3 (Z=1.3), the term ‘potential MFS’ should be proposed to use until the aorta reaches threshold. Herein, the second patient here should be categorized as ‘potential MFS’ (**Table.1**), with a total of 8 points for the systemic score according to the revised Ghent Criteria (LOEYS *et al*. 2010). Heterozygous *FBN1* missense variants c.4890_4891delGTinsTG (p.Gln1630_Cys1631delinsHisGly) were identified in the proband via targeted NGS and validated by Sanger sequencing (**Fig.1H**). The missense variants were confirmed to be *in cis* and comprising a complex allele resulting from a dinucleotide variant by visualizing read alignment views from the Integrative Genomics Viewer (IGV) (**Fig.1I**). Notably, the missense variant of c.4891T>G (p.Cys1631Gly) has been previously reported as a disease-causing mutation in a classical MFS individual, whose clinical course and phenotype was characterized by aortic root dissection, aortic root dilation, mitral valve regurgitation, tricuspid valve prolapse, *ectopia lentis* (ARBUSTINI *et al*. 2005). Conservation analysis of amino acid residues of Gln1630 and Cys1631in *FBN1* demonstrate that they are highly conserved throughout evolution and across many selected species, suggesting they are required for the normal function of the protein. Thus, the monoallelic missense variants were most likely damaging and putatively deleterious and possibly accounted for ‘potential MFS’ in this proband (**Fig.1J**).

#### Subject XH601

The proband was a 9-year-old Chinese girl (**Fig.1K, L**). When she was 4 years old at the time of her first evaluation, she presented with a height of 101 cm (P_25_; -1SD), an extremely thin body habitus with a weight of 12 kg (P_3_, -3SD). Physical examination revealed bilateral arachnodactyly (**Fig.1M, N**), uneven asymmetric back whilst bending and congenital dislocation of the hip joint. Whole-spine X ray examination displayed severe right thoracic scoliosis with a main Cobb angle of 117 degrees (**Fig.1O, P**). Transthoracic echocardiogram demonstrated anterior and posterior mitral leaflet prolapse and moderate mitral valve insufficiency without apparent aortic diameter enlargement at the sinuses of Valsalva or aortic root dissection. Ocular system abnormalities indicated bilateral down-slanting palpebral fissures, epicanthus and astigmatism. However, there were no signs or symptoms of ectopia lentis. The predominant features in this patient were an extreme congenital lack of subcutaneous fat, consistent with a lipodystrophy by physical exam, and a subsequent bodily progeroid appearance. However, this proband did not meet revised Ghent Criteria for classical MFS. Consecutive follow-ups and assessment of the proband were conducted. Upon genotype driven reverse phenotyping of the proband when she was 9, the proportion of upper and lower segment lengths was basically normal with a ratio of 0.87. The proband had a height of 136 cm (P_50_, 0). For comparison, her father’s height measurement was 170 cm and her mother’s was 160 cm. The body mass index (BMI) of the proband was extremely low (11.1 kg/m^2^) potentially due to disproportionate weight gain by age. More specifically, the patient had an extremely thin habitus with a weight of 20.5 Kg (P_3_, -3SD). Furthermore, hand and foot radiographs to potentially assess ‘bone age’ suggested bilateral metacarpophalangeal joint dislocation, interosseous atrophy and dolichostenomelia (**Fig.1Q, R**). The proband was eventually diagnosed with Marfanoid-progeroid-lipodystrophy syndrome (**Table.2**). Through targeted NGS, a heterozygous variant of c.8275_8291delGAGAAGACAGCCATCTT (c.8275_8291del; p.Glu2759Cysfs*9) in *FBN1* was identified in the proband. We consequently confirmed this variant in subject XH601 and it was indeed *de novo* through parental Sanger sequencing and trio analysis (**Fig.1S**). In addition to Sanger sequencing validation, we also performed cloning sequencing on Subject XH601 to further verify that this indel is subject to frame-shift variation (data not shown).

**Table 2.**
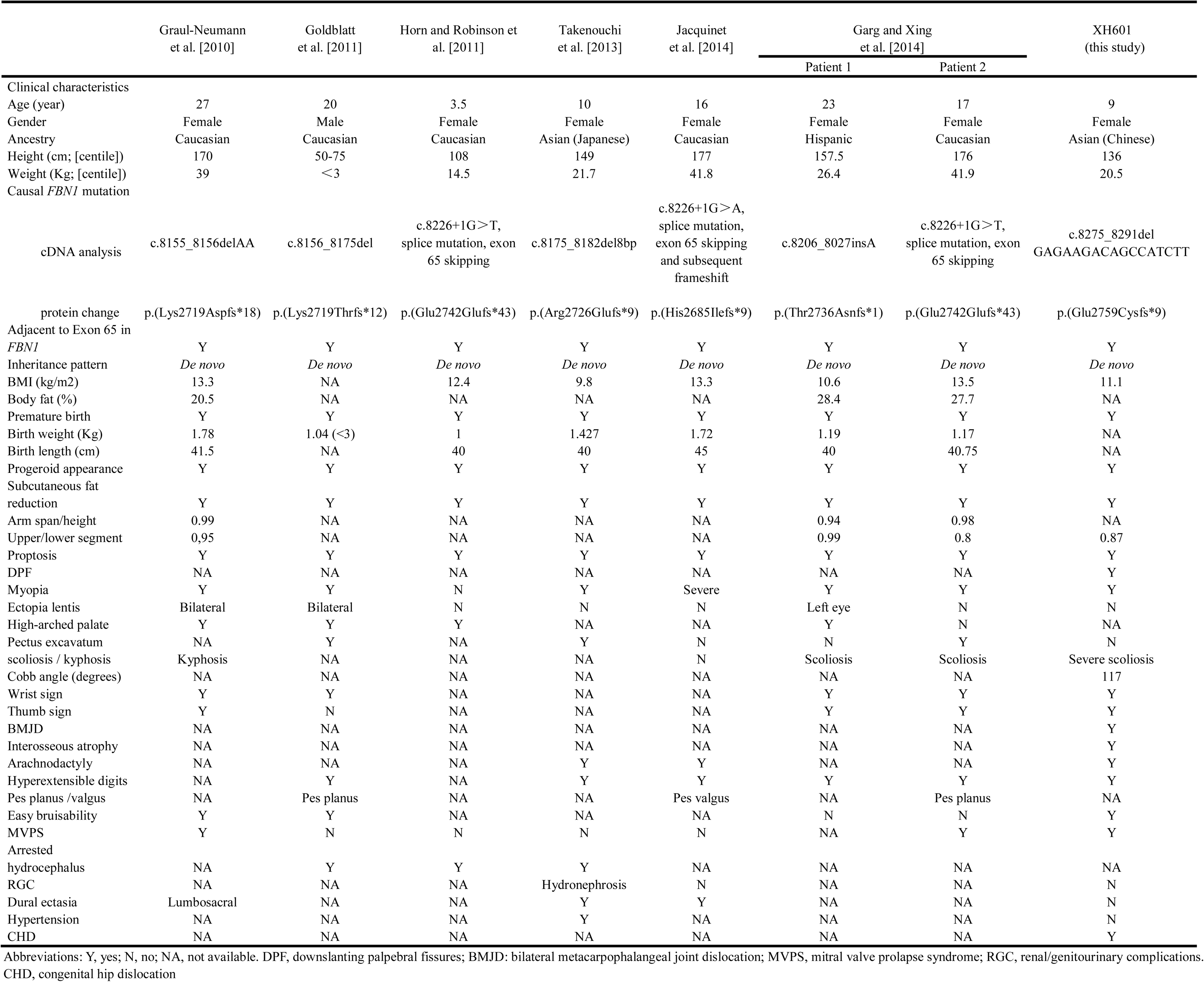
Clinical manifestations of previously reported and our patient XH601 with Marfanoid-progeroid-lipodystrophy (MPL) syndrome with allelic truncating mutations in *FBN1*

Moreover, these two allelic truncating variants of p.Tyr2596Thrfs*86 and p.Glu2759Cysfs*9 were not present in ExAC, 1000 Genome, ESP6500 database, universal mutation database (UMD) (COLLOD-BEROUD *et al*. 2003) and our in-house database (with 2000+ Chinese exomes). Clinical and genetic characteristics of subject XH253 with classical MFS and XH474 with ‘potential MFS’ based on the revised Ghent Criteria (LOEYS *et al*. 2010) are presented in Table 1.

### NMD-degradation prediction

The variant of Y2596Tfs*86 identified in XH253 was a single nucleotide frameshift deletion leading to a premature termination codon (PTC) in Exon 64, while E2759Cfs*9 identified in XH601 was a frameshift deletion leading to a PTC in Exon 66, the final exon of *FBN1* (**Fig. 2A**). Although both variants are very close in linear space, they may lead to distinct alterations to the *FBN1* transcript. To investigate this hypothesis, we performed NMD predictions of the two truncating variants on NMDescPredictor (COBAN-AKDEMIR *et al*. 2018). The result showed that Y2596Tfs*86 is predicted to be degraded by NMD, i.e. NMD^+^, while E2759Cfs*9 is predicted to escape NMD, i.e. NMD^-^ (**Fig. 2B**). Using the same analytical tools, we further tested whether previously reported protein-truncating *FBN1* frameshift variants implicated in MPLS subjects (GRAUL-NEUMANN *et al*. 2010; GOLDBLATT *et al*. 2011; TAKENOUCHI *et al*. 2013; GARG AND XING 2014) uniformly shared the mechanism of escaping NMD. Our analysis shows that all of the truncating *FBN1* frameshift variants within the C-terminal gene region may escape NMD, supporting the contention that the underlying disease mechanism in MPLS is the same and distinct from MFS (**Fig. 2B**). These findings implicate a mechanism distinct from loss-of-function (LoF) alleles, and potentially gain-of-function (GoF) variant alleles (COBAN-AKDEMIR *et al*. 2018), at the *FBN1* locus causing MPLS.

**Figure 2.**
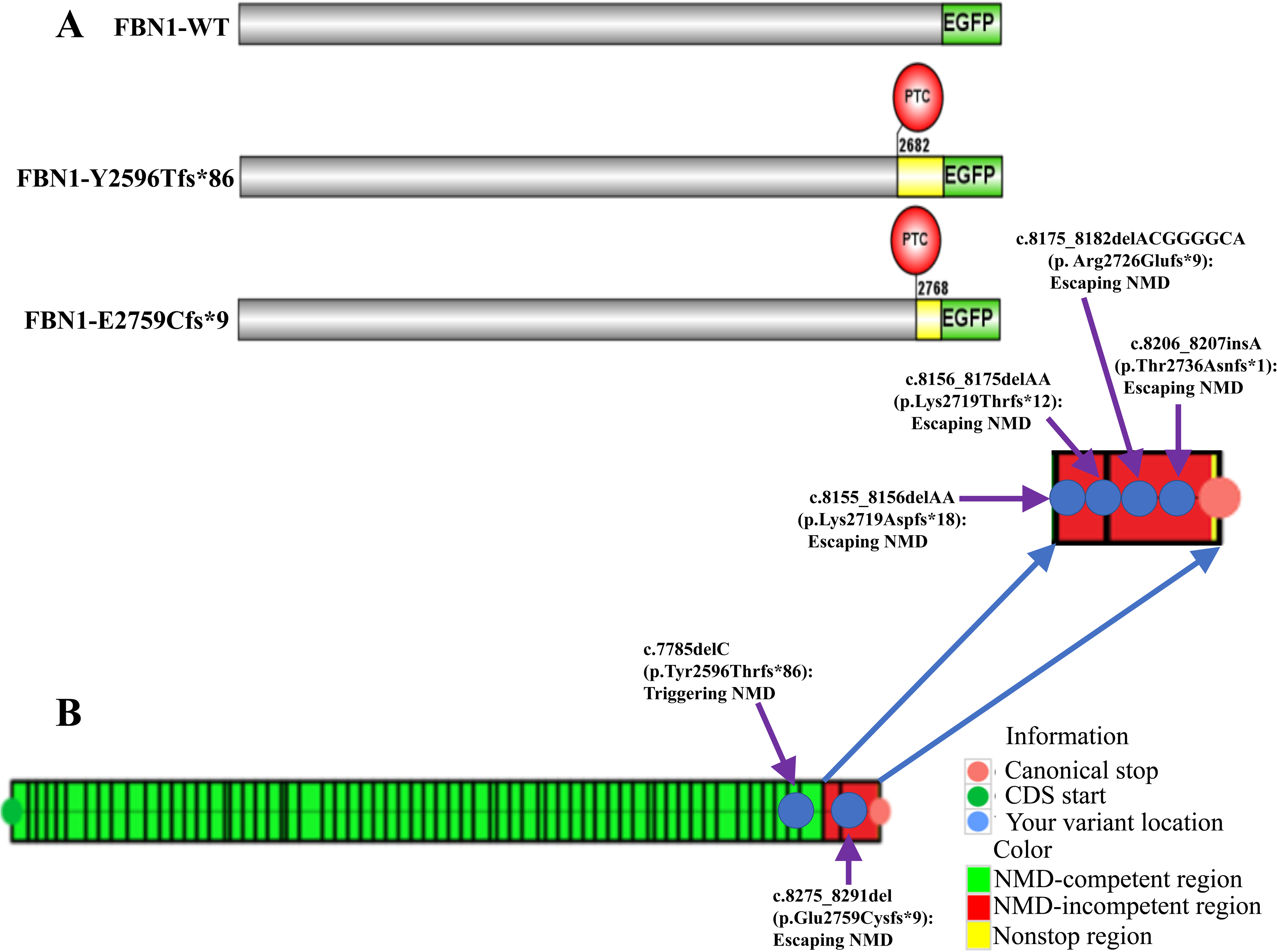
Schematic representations and NMD predictions of the p.Y2596Tfs*86 and p.E2759Cfs*9 mutations other MPLS-affected frameshift mutations in *FBN1* (A) Schematic diagrams of full-length WT and mutant EGFP-FBN1 plasmids transiently expressed in HEK293T. The numbers indicate the amino acid positions in *FBN1*. PTC location is marked with red circle. The resulting frameshift and reduction in the deduced amino acid sequence caused by the p.Y2596Tfs*86 and p.E2759Cfs*9 mutations are indicated by the yellow region, respectively. (B) Prediction of frameshift variants that are potentially subject to nonsense-mediated mRNA decay (NMD)-escape or NMD-degradation. The mutation of p.Y2596Tfs*86 in *FBN1* gene is predicted to be subject to degradation by triggering NMD. The p.E2759Cfs*9 mutation in *FBN1* gene is predicted to escape NMD. All of other MPLS-affected frameshift mutations in *FBN1* are predicted to escape NMD.

### Perturbation of native aggregation process by MPLS-causative truncating mutation

To further determine if the mutant FBN1 protein in MPLS is able to lead to dominant negative or GoF effects, plasmids expressing mutants (pEGFP-FBN1-Tyr2596Thrfs*86; pEGFP-FBN1-Glu2759Cysfs*9) and WT plasmid (pEGFP-FBN1) were co-transfected into HEK293T cells at a 1:1 ratio. After 48h of expression, lysates from cells expressing EGFP-FBN1 fusions were analyzed by SDD-AGE. Detection of SDS-resistant aggregates by SDD-AGE in cell lysates of HEK293T transiently expressing EGFP-FBN1 fusions were investigated with SDD-AGE and Western blot. Protein expression was induced for 48 h and detected with a monoclonal FBN1-specific antibody. Remarkably, WT plasmid (EGFP-FBN1 fusions) had the SDS-resistance properties of an amyloidogenic protein structure. Since amyloid formation is time- and concentration-dependent, we set the time of transient expression to 48 h. WT plasmid (EGFP-FBN1 fusions) were aggregated after 48 h expression, whereas co-transfection of pEGFP-FBN1-Glu2759Cysfs*9 and WT plasmid resulted in the presence of a small fraction of monomer protein. This suggests that pEGFP-FBN1-Glu2759Cysfs*9 apparently prohibited the native aggregation process of WT plasmid, possibly revealing an intracellular dominant-negative mechanism. In addition, we observed that co-transfection of pEGFP-FBN1-Tyr2596Thrfs*86 and WT plasmid presented a small fraction of aggregated protein. As this fraction was less than that of the WT plasmid, it can be suggested that Tyr2596Thrfs*86 may undergo an NMD mechanism thus leading to *FBN1* haploinsufficiency (**Fig.3A**). Moreover, these variations in the range of particle sizes were observed, and again were reproducible for individual EGFP-FBN1 fusions validated by another two independent replication experiments.

**Figure 3.**
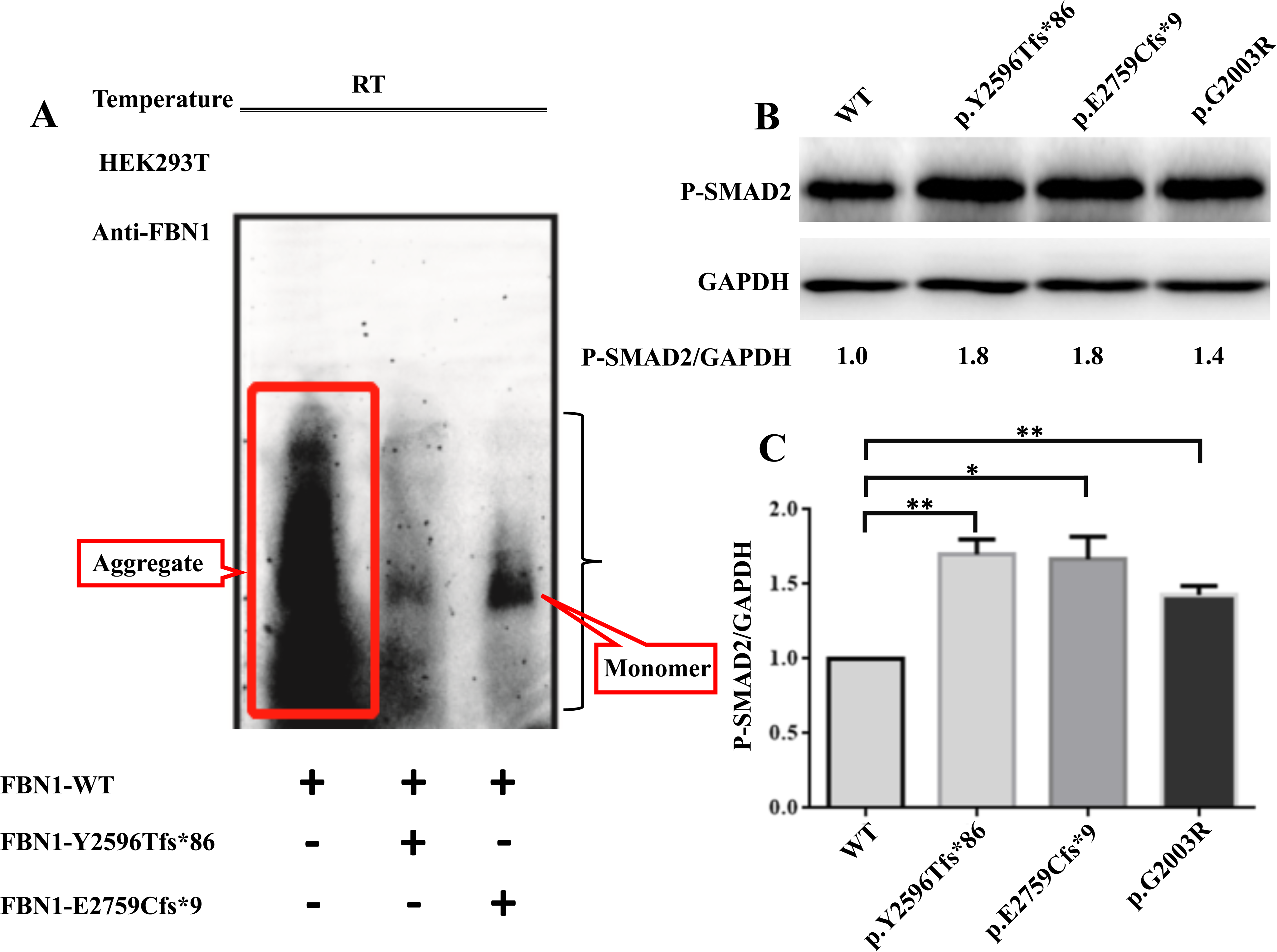
Functional effects of the Y2596Tfs*86 and E2759Cfs*9 mutations on *FBN1* protein expression (A) Detection of SDS-resistant aggregates by SDD-AGE in cell lysates of HEK293T transiently expressing EGFP-FBN1 fusions were investigated by SDD-AGE and Western blot. Expression of the proteins was induced for 48 h and detected with a monoclonal FBN1-specific antibody. (B) Elevated phosphorylation of SMAD2 (pSMAD2) in individuals with truncating *FBN1* variants, respectively. G2003R is used as a positive control. GAPDH is used as a loading control. pSMAD2/GAPDH ratio is shown normalized to the unaffected control (WT). (C) Gray-scale analysis results show significantly upregulated (pSMAD2) in constructs encoding truncating *FBN1* variants. EV denotes empty vector (pEGFP). RT: room temperature. Data are represented as mean ± SD of three independent experiments. * denotes P value <0.05 and ** denotes P value <0.01.

### Disruption of downstream TGF-**β** signaling

Previous studies have demonstrated that deleterious *FBN1* mutations causing Marfan syndrome result in upregulated endogenous transforming growth factor β (TGF-β) receptor signaling (ANDELFINGER *et al*. 2016). Such signaling can be measured in plasma or indirectly measured through aberrant activation of downstream targets, including excessive phosphorylation of SMAD2 (pSMAD2) (VERSTRAETEN *et al*. 2016). To determine the functional consequences of the novel truncating *FBN1* variant of E2759Cfs*9 in the MLPS patient, plasmids expressing mutant EGFP-FBN1 cDNAs were transfected into HEK293T cells. Transfected cells with truncating *FBN1* variants of Y2596Tfs*86 (P=0.007), E2759Cfs*9 (P=0.017) and Gly2003Arg (P=0.006) showed significantly elevated phosphorylation of SMAD2 (pSMAD2) in Western blots compared to the WT plasmid (**Fig.3B, C**). This observation is consistent with the contention that the TGF-β signaling pathway is perturbed in a SMAD-dependent manner in all 3 subjects.

### Genotypic and phenotypic features of reported MPLS patients

Intriguingly, the mechanism underlying *FBN1*-related diseases involves either loss-of-function (LoF) or NMD surveillance pathway escape as conveyed by allelic truncating mutations. Such distinct mechanisms could contribute to distinct disease phenotypes of varying severity. According to a previous report, *FBN1* LOF mutations led to classical MFS (PARK *et al*. 2017), while predicted NMD surveillance pathway escape in *FBN1* can cause a MPLS phenotype (ROMERE *et al*. 2016), Patient XH601 in our cohort harbored a heterozygous *de novo* variant of E2759Cfs*9 in *FBN1,* which is located in the final exon. Notably, our MPLS patients presented with bilateral down-slanting palpebral fissures, epicanthus and eye astigmatism. The main clinical features of this complicated disorder, clustered in Table 2, include accelerated aging and postnatal lipodystrophy, poor weight gain since birth, premature birth, hyperextensible digits and generalized subcutaneous fat reduction leading to a progeroid appearance of the body in all patients. Mental and motor development remain mostly normal. Overlapping clinical features of MPLS with MFS are phenotypically diverse. Ocular system involvement like myopia have been observed in all individuals, although hyperextensible joints, arachnodactyly, and other significant signs of classical MFS are not always present, i.e. mitral valve prolapse in 4/8, lumbosacral dural ectasia in 2/3 (5 data points unavailable), *pectus excavatum* in 3/8, and ectopia lentis in 3/8. Scoliosis was reported in two patients aged 23 and 17 years, and kyphosis was reported in a patient aged 27 years, but not described in the others (**Table.2**). Remarkably, the major Cobb angles of Subject XH253 with classical MFS and Subject XH474 with MASS were 43 degrees and 85 degrees, respectively. Our MPLS patient presented a thoracic scoliosis with a major Cobb angle magnitude of 117 degrees—a condition much more severe compared to the MFS patients, counterparts.

## Discussion

In the present report, we provide direct evidence for potential dominant negative alleles in MPLS patients caused by truncating *FBN1* variants escaping NMD. These genetic and functional investigations also include description of the first patient with MPLS of Chinese ancestry.

Several case reports previously identified an association between monogenic *FBN1* mutations and reduction of subcutaneous fat and/or progeroid features (GRAUL-NEUMANN *et al*. 2010; GOLDBLATT *et al*. 2011; TAKENOUCHI *et al*. 2013; JACQUINET *et al*. 2014; PASSARGE *et al*. 2016). Here, we have provided further evidence for the existence of a distinct genetic disease, MPLS, caused by *FBN1* mutations. The MPLS patient in our study is characterized by accelerated aging and postnatal lipodystrophy, disproportionate weight gain since birth, severe scoliosis, downslanting palpebral fissures, bilateral metacarpophalangeal joint dislocation, mitral valve prolapse, bilateral interosseous atrophy, and congenital dislocation of the hip. Intriguingly, this patient was previously diagnosed with Marfanoid disease (**Table.1**) due to an unclear molecular diagnosis. We consequently performed targeted NGS on the affected proband and her unaffected parents. As a result, we identified c.8275_8291del(p.Glu2759Cysfs*9) in *FBN1* as the responsible variant for the proband. Sanger sequencing was then conducted on genomic DNA from both parents to confirm the *de novo* occurrence and segregation with phenotypes. Subsequently, we proceeded with a comprehensive phenotypic analysis of all family members through intensive clinical follow-up. By combining genetic data with reverse phenotyping, we eventually diagnosed the proband with MPLS. Notably, the severity of spinal deformity in the MPLS individual was significantly more severe than that in Subject XH253 and XH474 (with classical MFS and potential MFS, respectively). We consider that MPLS is a distinct fibrillinopathy because it can be clinically distinguished from other fibrillinopathies, including classical MFS. To our knowledge, there are no other syndromes that simultaneously comprise the clinical phenotypes of marfanoid features, accelerated aging, postnatal lipodystrophy, poor weight gain since birth, and progeroid appearance. Particularly, the involvement of both reduced subcutaneous fat tissue and progeroid appearance is a prominent characteristic of MPLS. These comprehensive findings predominate in determining the severity of defects in individuals and thus significantly influence the prognosis: lipodystrophic disorders are frequently associated with metabolic disturbance, such as insulin resistance and life-threatening hypertriglyceridemia (VANTYGHEM *et al*. 2012). Such potential metabolic disturbances are particularly important to note prior to surgical intervention as has been noted for the ocular-scoliotic form of Ehlers Danlos syndrome due to lysl hydroyase deficiency (YEOWELL *et al*. 1995). Secondly, elucidation of the phenotype of MPLS has been limited due to the paucity of prior clinical reports, with only seven relevant patients previously described. Up to now, it is known only to be caused by a single gene with a monogenic AD inheritance pattern for the disease trait. Finally, we have shown that, in terms of pathogenic mechanisms, the truncating *FBN1* mutations that cause MPLS cluster in 3’ gene regions encoding the extreme C-terminal domains and these variant alleles represent a subset that differ from those that cause classical MFS. Most of such mutations will be anticipated to escape NMD and we propose these to function via a potential mechanism between GoF and dominant negative and not a LoF mechanism.

Fibrillin-1 acts as the precursor to a recently-described glucogenic hormone: aprosin (DAVIS *et al*. 2016a). Specifically, Exon 65 of *FBN1* encodes 11 amino acids, while Exon 66 of *FBN1* encodes 129 amino acids of asprosin (ROMERE *et al*. 2016). Recently, Chen et.al. have successfully constructed a gene-edited rabbit model with a truncated C-terminus of fibrillin-1 involving the ultimate two exons which could recapitulate the histopathological alterations and functional defects associated with MPLS (CHEN *et al*. 2018). Since individuals with low fibrillin-1 level may fail to differentiate adipocytes and/or to accumulate adipocyte lipids, truncating variants located adjacent to the two exons of *FBN1* are prone to cause lipodystrophic phenotypes (DAVIS *et al*. 2016b).

PTCs located in the final coding exon are specifically prone to escape NMD and so are processed distinctly from those in internal exons in terms of transcript degradation, leading to the stable translation of truncated proteins (BAYRAM *et al*. 2017). Excessive amounts of mutated protein are therefore produced (INOUE *et al*. 2004), leading to a dominant negative molecular mechanism and the severe MPLS disease phenotype. Consistent with this interpretation, all truncating mutations associated with MPLS are located adjacent to the penultimate exon (Exon 65) and are predicted to escape NMD (**Figure.4**). From this perspective, it is perhaps most parsimonious to postulate that the incorporation of relatively few mutant monomers would be sufficient to impair the systemic processes of microfibrillar assembly and function.

**Figure 4.**
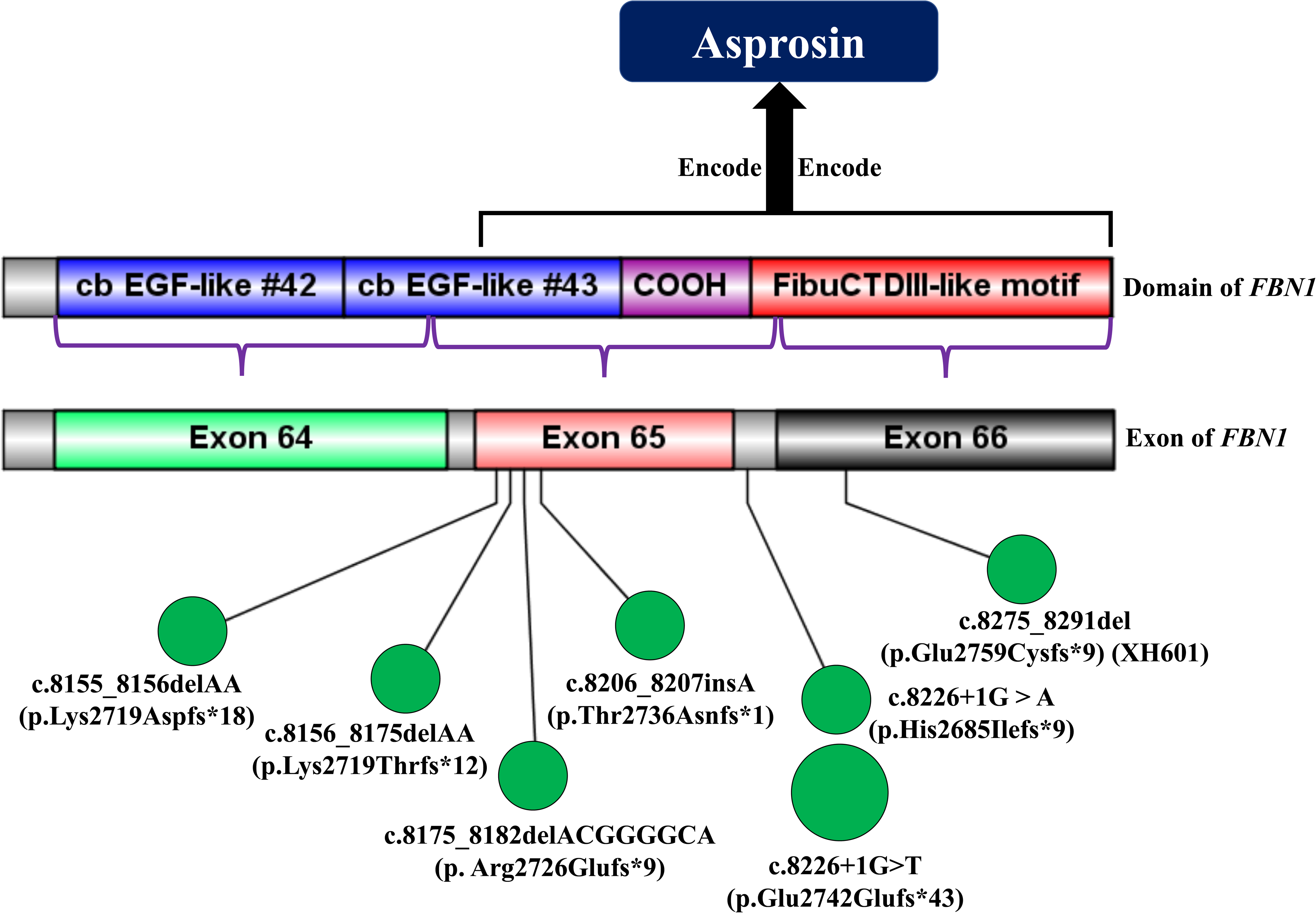
The partial gene structure of the C-terminus of *FBN1* and overview of previously reported truncating variants in and around Exon 65 in patients presenting with MPLS Blue, purple and red boxes denote specific domain. Light green, orange and black boxes denote exons and the gray area between the two boxes denote introns. All truncating variants were annotated by small green circles except that c.8226+1G>T previously reported twice was annotated by a big circle. COOH denotes COOH unique region. Deep blue box denotes purported asprosin encoded by the exons of *FBN1*.

Aberrant activation of the TGF-β pathway has been observed in MFS and may account for some of the musculoskeletal deformities, such as scoliosis (BUCHAN *et al*. 2014). The MPLS patient with a truncating *FBN1* variant presented with severe scoliosis, which suggests that the protein change also alters TGF-β signaling. Our results showed upregulation of the TGF-β pathway in plasmids expressing mutant EGFP-FBN1 cDNAs, confirming that the variant of E2759Cfs*9 identified in the MPLS patient has functional effects in a SMAD-dependent manner in all 3 subjects.

Our study indicated that the two phenotypes associated with *FBN1* mutations, MPLS and MFS are caused by two distinct molecular mechanisms. The more severe phenotype of MPLS, which had its first report in the Chinese population, is caused by nonsense mutations that produce truncated *FBN1* mutant proteins. These proteins disrupt the process of native amyloid-like structure conformation and exert potent intracellular dominant-negative activity. In this view, abnormal protein derived from the mutant allele interacts and interferes with protein derived from the normal allele, consequently resulting in substantial loss of function. Supportive evidence includes (a) *de novo* monogenic AD inheritance pattern, and (b) aggregation of fibrillin-1 in which the mutated monomer derived from the mutant allele impairs the fundamental structure of its wildtype protein derived from the normal allele counterpart. This deficiency could be supported by SDD-AGE analysis, suggesting that dominant negative interference is not restricted to enhanced proteolytic clearance of mutant microfibrils over time (BRENN *et al*. 1996), but rather could also occur at the level of intracellular aggregate formation processes. The more moderate phenotype, classical MFS, is caused by nonsense mutations that activate the NMD RNA surveillance pathway, thereby degrading mutant transcripts and resulting in *FBN1* haploinsufficiency. The data support the concept of distinct pathogenetic mechanisms, GoF *versus* LoF (BAYRAM *et al*. 2017; COBAN-AKDEMIR *et al*. 2018; POLI *et al*. 2018), for each well-established subgroup of fibrillinopathy, which mostly hinges on the intrinsic features of fibrillin-1 mutations. It is of crucial significance to pinpoint and appreciate the locus heterogeneity, clinical relevance and overlapping clinical features with other syndromes/disorders in a phenotype-oriented approach that characterizes much of clinical medicine practice (WHITE *et al*. 2018).

In conclusion, we provide direct evidence of dominant negative effects of truncating *FBN1* variants predicted to escape NMD in MPLS patients. Our study expands the mutational spectrum of *FBN1* and highlights the potential molecular mechanism for MPLS patients, which facilitates our understanding of genotype-phenotype correlations in terms of *FBN1* to provide effective genetic counseling, implementation and timing of therapy (e.g. mitigation of TGF-β hypersignaling, surgical intervention for cardiovascular complications or for scoliosis), or early intervention.

## Acknowledgments

We are grateful to the patients, their families, and genetic counselors for providing samples and clinical histories. This research was funded in part by the National Natural Science Foundation of China (81672123 to JG.Z, 81822030 to N.W., 81772299 to ZH.W., 81772301 to GX.Q.,), Beijing Natural Science Foundation (7172175 to N.W.; 7184232 for S.L., 5184037 to TS.S.), CAMS Initiative for Innovative Medicine (2016-I2M-3-003 to GX.Q. and N.W., 2016-I2M-2-006 and 2017-I2M-2-001 to ZH.W.), the Central Level Public Interest Program for Scientific Research Institute (2018RC31003 to N.W. and M.L.), the National Key Research and Development Program of China (No. 2018YFC0910506 to N.W. and ZH.W., 2016YFC0901501 to SY.Z.), the US National Institutes of Health, National Institute of Neurological Disorders and Stroke (NINDS R01 NS058529 and R35 NS105078 to J.R.L.), National Human Genome Research Institute/National Heart, Lung, and Blood Institute (NHGRI/NHLBI UM1 HG006542 to J.R.L.), and the National Human Genome Research Institute (NHGRI K08 HG008986 to J.E.P.).

## References

Andelfinger, G., B. Loeys and H. Dietz, 2016 A Decade of Discovery in the Genetic Understanding of Thoracic Aortic Disease. Can J Cardiol 32: 13–25.

Arbustini, E., M. Grasso, S. Ansaldi, C. Malattia, A. Pilotto et al., 2005 Identification of sixty-two novel and twelve known *FBN1* mutations in eighty-one unrelated probands with Marfan syndrome and other fibrillinopathies. Hum Mutat 26: 494.

Asan, Y. Xu, H. Jiang, C. Tyler-Smith, Y. Xue et al., 2011 Comprehensive comparison of three commercial human whole-exome capture platforms. Genome Biol 12: R95.

Bayram, Y., J. J. White, N. Elcioglu, M. T. Cho, N. Zadeh et al., 2017 REST Final-Exon-Truncating Mutations Cause Hereditary Gingival Fibromatosis. Am J Hum Genet 101: 149–156.

Berchowitz, L. E., G. Kabachinski, M. R. Walker, T. M. Carlile, W. V. Gilbert et al., 2015 Regulated Formation of an Amyloid-like Translational Repressor Governs Gametogenesis. Cell 163: 406–418.

Brenn, T., T. Aoyama, U. Francke and H. Furthmayr, 1996 Dermal fibroblast culture as a model system for studies of fibrillin assembly and pathogenetic mechanisms: defects in distinct groups of individuals with Marfan’s syndrome. Lab Invest 75: 389–402.

Buchan, J. G., D. M. Alvarado, G. E. Haller, C. Cruchaga, M. B. Harms et al., 2014 Rare variants in *FBN1* and *FBN2* are associated with severe adolescent idiopathic scoliosis. Hum Mol Genet 23: 5271–5282.

Chen, M., B. Yao, Q. Yang, J. Deng, Y. Song et al., 2018 Truncated C-terminus of fibrillin-1 induces Marfanoid-progeroid-lipodystrophy (MPL) syndrome in rabbit. Dis Model Mech 11.

Coban-Akdemir, Z., J. J. White, X. Song, S. N. Jhangiani, J. M. Fatih et al., 2018 Identifying Genes Whose Mutant Transcripts Cause Dominant Disease Traits by Potential Gain-of-Function Alleles. Am J Hum Genet 103: 171–187.

Collod-Beroud, G., S. Le Bourdelles, L. Ades, L. Ala-Kokko, P. Booms et al., 2003 Update of the UMD-FBN1 mutation database and creation of an *FBN1* polymorphism database. Hum Mutat 22: 199–208.

Davis, M. R., E. Arner, C. R. Duffy, P. A. De Sousa, I. Dahlman et al., 2016a Datasets of genes coexpressed with *FBN1* in mouse adipose tissue and during human adipogenesis. Data Brief 8: 851–857.

Davis, M. R., E. Arner, C. R. Duffy, P. A. De Sousa, I. Dahlman et al., 2016b Expression of *FBN1* during adipogenesis: Relevance to the lipodystrophy phenotype in Marfan syndrome and related conditions. Mol Genet Metab 119: 174–185.

Dietz, H. C., 2015 Potential Phenotype-Genotype Correlation in Marfan Syndrome: When Less is More? Circ Cardiovasc Genet 8: 256–260.

Duerrschmid, C., Y. He, C. Wang, C. Li, J. C. Bournat et al., 2017 Asprosin is a centrally acting orexigenic hormone. Nat Med 23: 1444–1453.

Garg, A., and C. Xing, 2014 De novo heterozygous *FBN1* mutations in the extreme C-terminal region cause progeroid fibrillinopathy. Am J Med Genet A 164a: 1341–1345.

Goldblatt, J., J. Hyatt, C. Edwards and I. Walpole, 2011 Further evidence for a marfanoid syndrome with neonatal progeroid features and severe generalized lipodystrophy due to frameshift mutations near the 3’ end of the *FBN1* gene. Am J Med Genet A 155a: 717–720.

Graul-Neumann, L. M., T. Kienitz, P. N. Robinson, S. Baasanjav, B. Karow et al., 2010 Marfan syndrome with neonatal progeroid syndrome-like lipodystrophy associated with a novel frameshift mutation at the 3’ terminus of the FBN1-gene. Am J Med Genet A 152a: 2749–2755.

Inoue, K., M. Khajavi, T. Ohyama, S. Hirabayashi, J. Wilson et al., 2004 Molecular mechanism for distinct neurological phenotypes conveyed by allelic truncating mutations. Nat Genet 36: 361–369.

Jacquinet, A., A. Verloes, B. Callewaert, C. Coremans, P. Coucke et al., 2014 Neonatal progeroid variant of Marfan syndrome with congenital lipodystrophy results from mutations at the 3’ end of *FBN1* gene. Eur J Med Genet 57: 230–234.

Loeys, B. L., H. C. Dietz, A. C. Braverman, B. L. Callewaert, J. De Backer et al., 2010 The revised Ghent nosology for the Marfan syndrome. J Med Genet 47: 476–485.

Park, J. W., L. Yan, C. Stoddard, X. Wang, Z. Yue et al., 2017 Recapitulating and Correcting Marfan Syndrome in a Cellular Model. Int J Biol Sci 13: 588–603.

Passarge, E., P. N. Robinson and L. M. Graul-Neumann, 2016 Marfanoid-progeroid-lipodystrophy syndrome: a newly recognized fibrillinopathy. Eur J Hum Genet 24: 1244–1247.

Poli, M. C., F. Ebstein, S. K. Nicholas, M. M. de Guzman, L. R. Forbes et al., 2018 Heterozygous Truncating Variants in *POMP* Escape Nonsense-Mediated Decay and Cause a Unique Immune Dysregulatory Syndrome. Am J Hum Genet 102: 1126–1142.

Romere, C., C. Duerrschmid, J. Bournat, P. Constable, M. Jain et al., 2016 Asprosin, a Fasting-Induced Glucogenic Protein Hormone. Cell 165: 566–579.

Schneider, C. A., W. S. Rasband and K. W. Eliceiri, 2012 NIH Image to ImageJ: 25 years of image analysis. Nat Methods 9: 671–675.

Song, Y. H., G. H. Kim, H. W. Yoo and J. B. Kim, 2012 Novel de novo nonsense mutation of *FBN1* gene in a patient with Marfan syndrome. J Genet 91: 233–235.

Takenouchi, T., M. Hida, Y. Sakamoto, C. Torii, R. Kosaki et al., 2013 Severe congenital lipodystrophy and a progeroid appearance: Mutation in the penultimate exon of *FBN1* causing a recognizable phenotype. Am J Med Genet A 161a: 3057–3062.

Vantyghem, M. C., A. S. Balavoine, C. Douillard, F. Defrance, L. Dieudonne et al., 2012 How to diagnose a lipodystrophy syndrome. Ann Endocrinol (Paris) 73: 170–189.

Verstraeten, A., M. Alaerts, L. Van Laer and B. Loeys, 2016 Marfan Syndrome and Related Disorders: 25 Years of Gene Discovery. Hum Mutat 37: 524–531.

White, J. J., J. F. Mazzeu, Z. Coban-Akdemir, Y. Bayram, V. Bahrambeigi et al., 2018 WNT Signaling Perturbations Underlie the Genetic Heterogeneity of Robinow Syndrome. Am J Hum Genet 102: 27–43.

Wu, N., X. Ming, J. Xiao, Z. Wu, X. Chen et al., 2015 *TBX6* null variants and a common hypomorphic allele in congenital scoliosis. N Engl J Med 372: 341–350.

Yeowell, H. N., L. C. Walker, M. K. Marshall, S. Murad and S. R. Pinnell, 1995 The mRNA and the activity of lysyl hydroxylase are up-regulated by the administration of ascorbate and hydralazine to human skin fibroblasts from a patient with Ehlers-Danlos syndrome type VI. Arch Biochem Biophys 321: 510–516.

